# Structural basis for gating inhibition by the cytoplasmic domain in HCN1 channels

**DOI:** 10.1101/2025.10.14.682313

**Authors:** V. Zorzini, J.M. Kleiz-Ferreira, M. Brams, R. Gallardo, S. De Gieter, J. Kusch, C. Ulens

## Abstract

Hyperpolarization-activated, cyclic nucleotide-gated (HCN) channels generate rhythmic electrical activity in cardiac and neuronal tissues, with isoform-specific cAMP sensitivity remaining poorly understood. While HCN2 exhibits strong cAMP regulation, HCN1 shows minimal response. To investigate the structural basis of this divergence, we analyzed two engineered HCN1 variants using cryo-electron microscopy. One variant (HCN112) incorporates the C-linker and CNBD from HCN2 into the HCN1 backbone and exhibited enhanced cAMP sensitivity, with structural analysis revealing a compressed cytoplasmic domain arrangement that may facilitate regulatory interactions. In contrast, the truncated HCN1ΔC variant (lacking the cytoplasmic domain) displayed an intermediately open pore conformation, supporting auto-inhibitory regulation by the CNBD in wild-type channels. These structural insights elucidate how domain-specific interactions modulate cAMP-dependent gating and intrinsic auto-inhibition, resolving long-standing questions about mechanistic divergence among HCN isoforms. Our findings not only shed new light on the structural mechanisms underlying isoform-specific cAMP sensitivity but also have implications for the development of therapeutic strategies targeting HCN channels in neurological and cardiac disorders.

**SIGNIFICANCE STATEMENT:** Hyperpolarization-activated cyclic nucleotide-gated (HCN) channels are essential regulators of rhythmic electrical activity in the heart and brain, and their dysfunction is linked to disorders such as epilepsy, depression, and cardiac arrhythmias. Although HCN channel isoforms display highly divergent responses to cAMP modulation, the structural basis for these differences has remained elusive. This study reveals, through cryo-EM structures, how domain-specific interactions within the cytoplasmic regions of HCN1 channels underlie their unique gating and auto-inhibition properties, as well as their muted cAMP sensitivity compared to HCN2. These insights resolve long-standing mechanistic questions about HCN channel regulation and pave the way for rational design of targeted therapies that selectively modulate HCN isoform activity in neurological and cardiac disease.

## INTRODUCTION

Ion channels are integral membrane proteins that act as molecular switches of electrochemical signaling in the central nervous system. More specifically, the Hyperpolarization-activated Cyclic Nucleotide-sensitive (HCN) channels belong to a family of cation channels that is activated by hyperpolarizing potentials and regulated by cyclic AMP (cAMP). HCN channels serve as nonselective voltage-gated cation channels that contribute to spontaneous rhythmic activity in the plasma membranes of heart and brain cells. This type of channel is also concentrated in cortical and hippocampal pyramidal cell dendrites, where it plays an important role in determining synaptic input integration and thus neuronal output. Therefore, it has been suggested to be involved in physiological processes such as cognition as well as pathophysiological states such as epilepsy. Recent evidence suggests that these channels may also be therapeutic targets for treatment of depressive disorders^1^.

Wang et al., 2001 have identified structural domains in HCN1 that are important for the basal voltage dependence of gating and the extent to which cAMP modulates the gating^2^. Specifically, channels containing either the core transmembrane domain or C-terminus of HCN2 channels activate at more negative voltages compared with channels that contain the corresponding regions of HCN1. Differences in cAMP modulation are largely localized to the C-terminus, with channels containing the HCN2 C-terminus showing a larger shift in response to cAMP compared to channels containing the HCN1 C-terminus^2^. Thus, removal of the entire C-terminus fully eliminates the response to cAMP. Yet the precise nature and molecular mechanism of this structural modulation is unknown.

Although the HCN1 structures in its apo-state and also cAMP-bound have already been solved by cryo-EM^3^, together with other HCN1 variants^4^, none of these structures showed an open gate and therefore they do not fully explain the HCN1 mechanism of gating. More recently, another group reports a full panorama of structures on HCN4, in its apo-state and cAMP-bound, in presence of different detergents. The group also presents a HCN4 structure with the gate open shedding light onto gating movement and ion permeation in HCN4 pacemaker channels^5^. Multiple other studies support the HCN role also in the function of sensory neurons and pain sensation, particularly pain associated with nerve or tissue injury^6^ . Therefore, understanding the structural basis of the HCN channel regulation could offer unique insights into the molecular basis of its modulation and its druggability.

However, although lots of structures have already been solved, the regulation of HCN gating activity is not fully understood. How the different domains change conformation, communicate between each other’s, and also inhibit gating remains unknown.

This study aims to structurally unveil how the HCN gating mechanisms are modulated by the C-terminus in its apo-state and by cAMP ligand binding. To understand the role of the C-terminus in the modulation of gating induced by cAMP, we have expressed and purified two HCN variants: HCN112 and HCN1ΔC^2,7^ . HCN112 is a chimera between the cytoplasmic NH2 terminus and transmembrane domain of HCN1 and the cytoplasmic C-terminus of HCN2. On the other hand, HCN1ΔC has a deletion corresponding to the full cytoplasmic C-terminus.

These constructs allowed us to understand a long-standing question whether the C-terminal domain acts in an auto-inhibitory manner on the gate of the channel.

## RESULTS AND DISCUSSION

### C-terminal region of HCN2 shows a compressed conformation in an HCN1 channel background

In recent years, several research groups have published three-dimensional structures for three of the four HCN isoforms, namely HCN1, HCN3 and HCN4 channels^3–5,8,9^. For HCN2, however, only the three-dimensional structure of the isolated CL-CNBD has been elucidated so far^10,11^. To study the C-terminal region of the HCN2 channel not only as an isolated protein but in a full channel environment, we took advantage of the HCN112 chimera (Figure 1A). This chimera is composed of the N-terminal domain and transmembrane domain of HCN1 and the C-terminal domain of HCN2 channels^2^.

**Figure 1.**
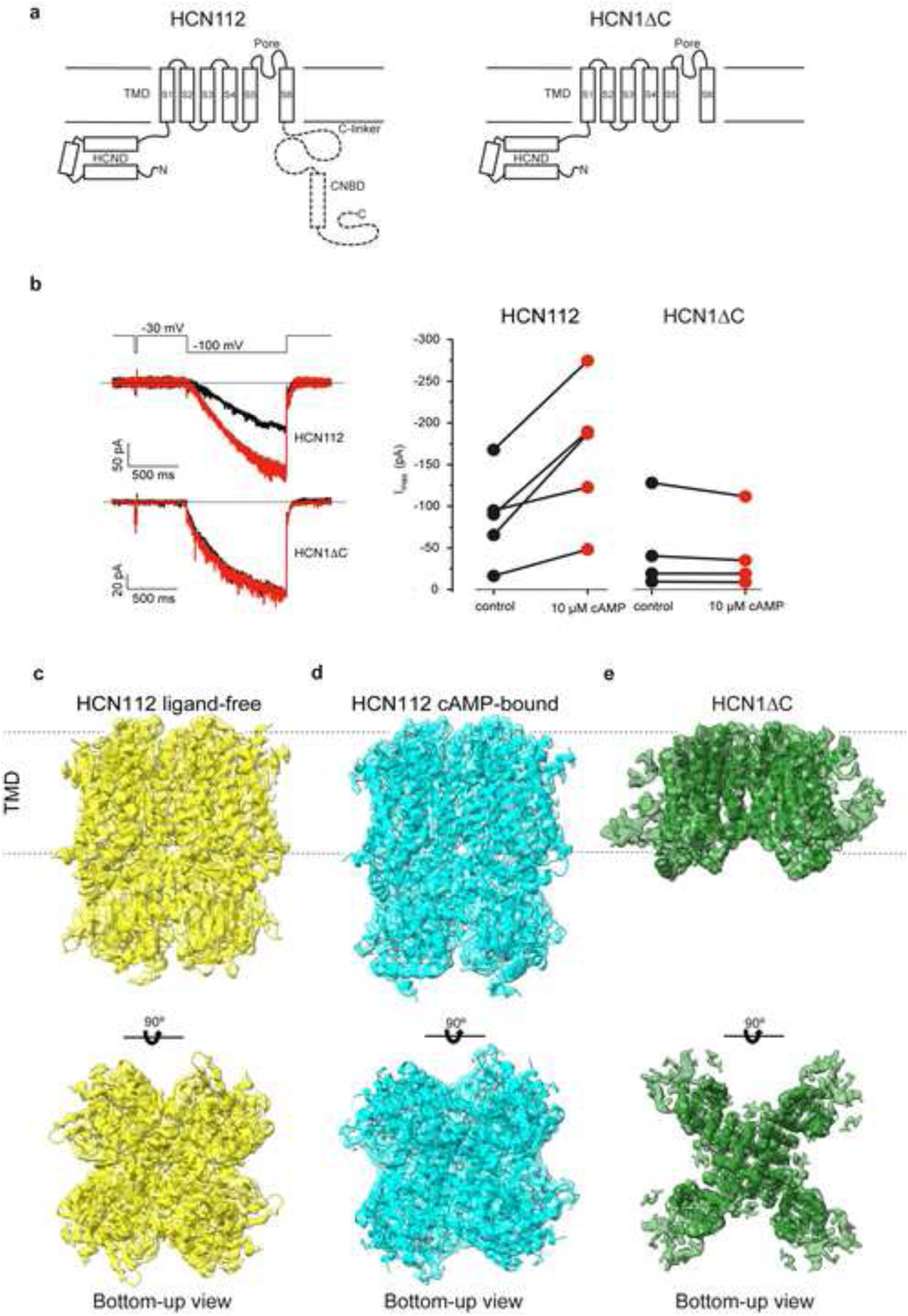
**Structure and function of HCN112 and HCN1**Δ**C channels. a**. Transmembrane topology of the HCN112 and HCN1ΔC channel constructs. The dashed line in HCN112 indicates the C-linker and CNBD, which have been replaced with the corresponding sequence from HCN2 channels. **b**. Representative current traces from HCN112 and HCN1ΔC channels expressed in *Xenopus* oocytes and evoked by a voltage step to −100 mV in the absence (black) and presence (red) of 10 μM cAMP. The graphs on the right show the summary of peak current amplitudes from 4 individual experiments. **c-e**. Cryo-EM maps of HCN112 ligand-free (yellow), HCN112 cAMP-bound (cyan) and HCN1ΔC (green). The 3D-structures are docked in the map and shown in cartoon representation. The top panels are side-views, the bottom panels are a bottom-up views.

To show that the CL-CNBD region in the HCN112 chimera is still functional, we tested the effect of cAMP on the current amplitude in inside-out macropatches of *Xenopus laevis* oocytes. Data presented in Figure 1B (left panel) show that the ability of cAMP to increase the maximum current amplitude, *I*_max_, evoked by a hyperpolarizing voltage jump is still intact after binding to an HCN2 CNBD in an HCN1 background. From this data we conclude that the HCN112 chimera is a valid tool to compare HCN1 and HCN2 CNBD structures in a functional channel system.

We expressed and purified HCN112 and HCN1ΔC according to Lee and MacKinnon^3^. Cryo-EM structures of HCN112 in the ligand-free and cAMP-bound state were determined to an overall resolution of 3.5 Å and 3.26 Å (Figure 1C-D). This is comparable to the resolution of HCN1 structures (3.5 Å) by Lee and MacKinnon^3^. The overall channel structure of HCN112 in the ligand-free state is comparable to HCN1 ligand-free (pdb code 5u6o) with an RMSD of 1.4250 Å. Similar to HCN1, HCN112 channels consist of four identical subunits forming a 4-fold symmetric channel which is surrounded by the membrane. It has a membrane-embedded set of six alpha-helices (S1-S6, transmembrane domain = TMD) constituting the pore (S5-S6) and the voltage sensor (S1-S4). The HCN112 C-terminal cytoplasmic domain is formed by the C-linker and the cyclic nucleotide binding domain (CNBD). Unique to HCN channels is the N-terminal HCN domain spanning the first 45 residues preceding S1 and forming a 3-alpha-helical domain. This domain is wedged between the voltage sensor and the cytoplasmic domain. The HCN domain contacts the S4 helix from the same subunit and the C-linker and CNBD from an adjacent subunit. This constrains the channel at the level of the C-linker disk^3^.

Besides these above-mentioned similarities in the overall channel structure, comparison of HCN112 and HCN1 in the ligand-free states reveals an important change in the quaternary conformation (Figure 2). In HCN112, the cytoplasmic domain and transmembrane domain are closer to each other in the direction perpendicular to the plane of the membrane, resulting in a ‘compressed’ mode of the TMD and CNBD (supplementary movie S1) in HCN112 compared to a ‘relaxed’ mode in HCN1. This compression amounts to ∼3.5 Å when measuring the distance from the top of the S5 helix (residue Q323) to the distal end of the CNBD (residue G547) (Figure 2A-C). Importantly, as a result of the compression movement in HCN112 we observe a drastic increase in the inter-chain interactions through hydrogen bonds and salt bridges (Figure 2D-F, Table 1). These interactions (indicated in purple, Figure 2E-F) are unique for HCN112 and not present in HCN1. For example, in the pore helix and middle of S6 there are unique H-bond interactions between chain A residues L357, Y386 and chain B residue C358. Additionally, at the top of the pore helix there is a unique H-bond between chain A residue A363 and chain B residue Q364. However, most of the unique interactions in HCN112 between chain A and B are localized in the C-linker, a region that has been shown repeatedly to be important for coupling ligand binding to channel opening^2,5,10,12–31^. In a recent study, the C-linker has also been implicated in the gating polarity of a CNBD channel chimera composed of the EAG pore domain in a HCN1 background^32^. The unique inter-chain salt bridges in the C-linker of HCN112 are D401-R297, R404-E283 and K430-E460. The unique inter-chain H-bonds are R405-Q398, Q408-S293, K412-S399/L400, Q416-K442, Y435-I449, Y439-E452 in the C-linker (Figure 2F). Lastly, there are unique inter-chain H-bond involving the CNBD, namely residues H517-F114 and R560-R112. As shown in Table 1 most of the H-bonds present in HCN112 but not in HCN1 are localized in the C-linker, a region important for coupling ligand binding to channel opening.

**Figure 2.**
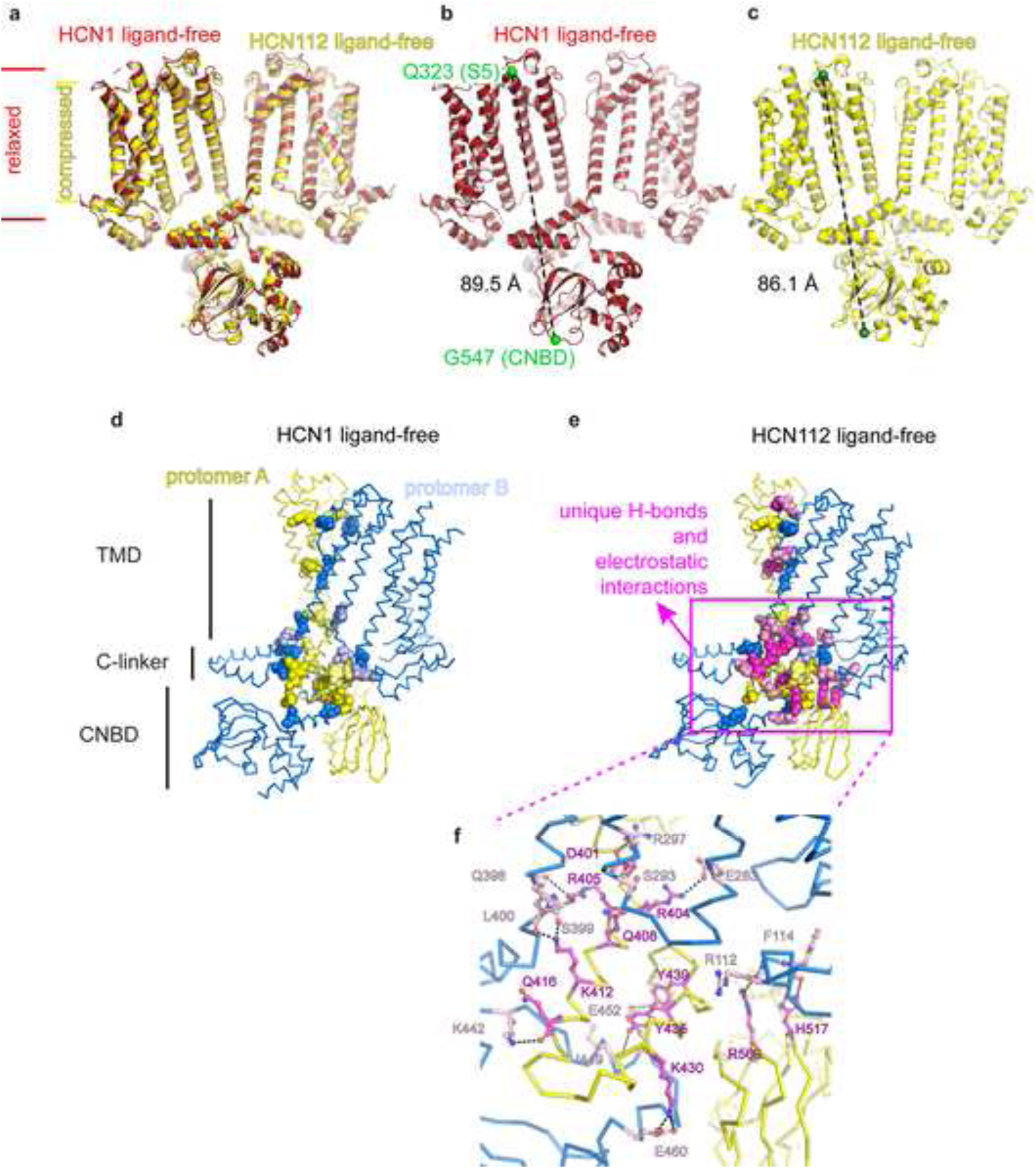
**HCN112 channels show a compressed conformation and unique inter-chain interactions. a-c**. Cartoon representations of HCN1 ligand-free (**b**, red, pdb code 5u6o), HCN112 ligand-free (**c**, yellow) and overlay (**a**). Only 2 opposing subunits are shown for clarity. The green spheres and the dashed line show the distance between the top of S5, residue Q323 and the distal end of the CNBD, residue G547. **d-e**. Inter-chain interactions between protomor A (yellow ribbon) and protomor B (blue ribbon) for HCN1 ligand-free (**d**, pdb code 5u6o) and HCN112 ligand-free (**e**). Interacting residues are shown in sphere representation. Hydrogen bonds are shown between yellow (protomor A) and dark blue (protomor B) residues. Salt bridges are shown between pale yellow (promotor A) and pale blue (protomor B) residues. Hydrogen bonds and salt bridges that are unique in HCN112 (**e**) are shown as magenta and pink residues. **f**. Detailed view of the uniquely interacting residues in the C-linker region. Residues of protomor A are shown in magenta ball and stick representation, residues of protomor B are shown in pink. Oxygen atoms are shown in red, nitrogen atoms are shown in blue.

**Table 1:** Unique H-bonds and salt bridges observed in HCN112 but not in HCN1.

| Salt bridges | cAMP-free | cAMP-bound |
| --- | --- | --- |
| CL | D401-R297 |  |
|  | R404-E283 |  |
|  | K430-E460 |  |
| **H-bonds** |  |  |
| TMD | L357-C358 |  |
|  | A363-Q364 |  |
|  | Y386-C358 |  |
| CL | R405-Q39 |  |
|  | Q408-S293 |  |
|  | K412-S399/L400 | K412-L400 |
|  | Q416-K442 |  |
|  |  | H421-N465 |
|  |  | Y434-E452 |
|  | Y435-I449 |  |
|  | Y439-E452 | Y439-E452 |
|  |  | V495-N454 |
|  |  | R548-E575 |
| CL-CNBD | H517-F114 |  |
|  | R560-R112 |  |

In summary, in HCN112 we observe a more compressed conformation of the structure in the plane perpendicular to the membrane compared to HCN1. As a result, there are more inter-chain interactions, especially in the C-linker which couples the CNBD to the TMD. We suggest that this compressed conformation is one of the reasons why cAMP works more efficiently in HCN2 than in HCN1 channels.

### cAMP binding breaks interactions in the compressed HCN112 conformation

To study the compressed mode and the unique inter-chain interactions in HCN112 CL-CNBD in more detail we next analyzed HCN112 in its cAMP-bound form.

The cryo-EM structure of cAMP-bound HCN112 was obtained from data at 3.26 Å resolution (Figure 3A-C). The structure of cAMP-bound HCN112 superimposes onto the ligand-free HCN112 structure with an RMSD of 0.542 Å. The largest conformational change upon cAMP-binding can be observed in the CNBD, which rotates and tilts away from the C-linker, and toward the C-terminal helices D and E (Supplementary movie S2). These helices rigidify and become more structured upon cAMP binding.

**Figure 3.**
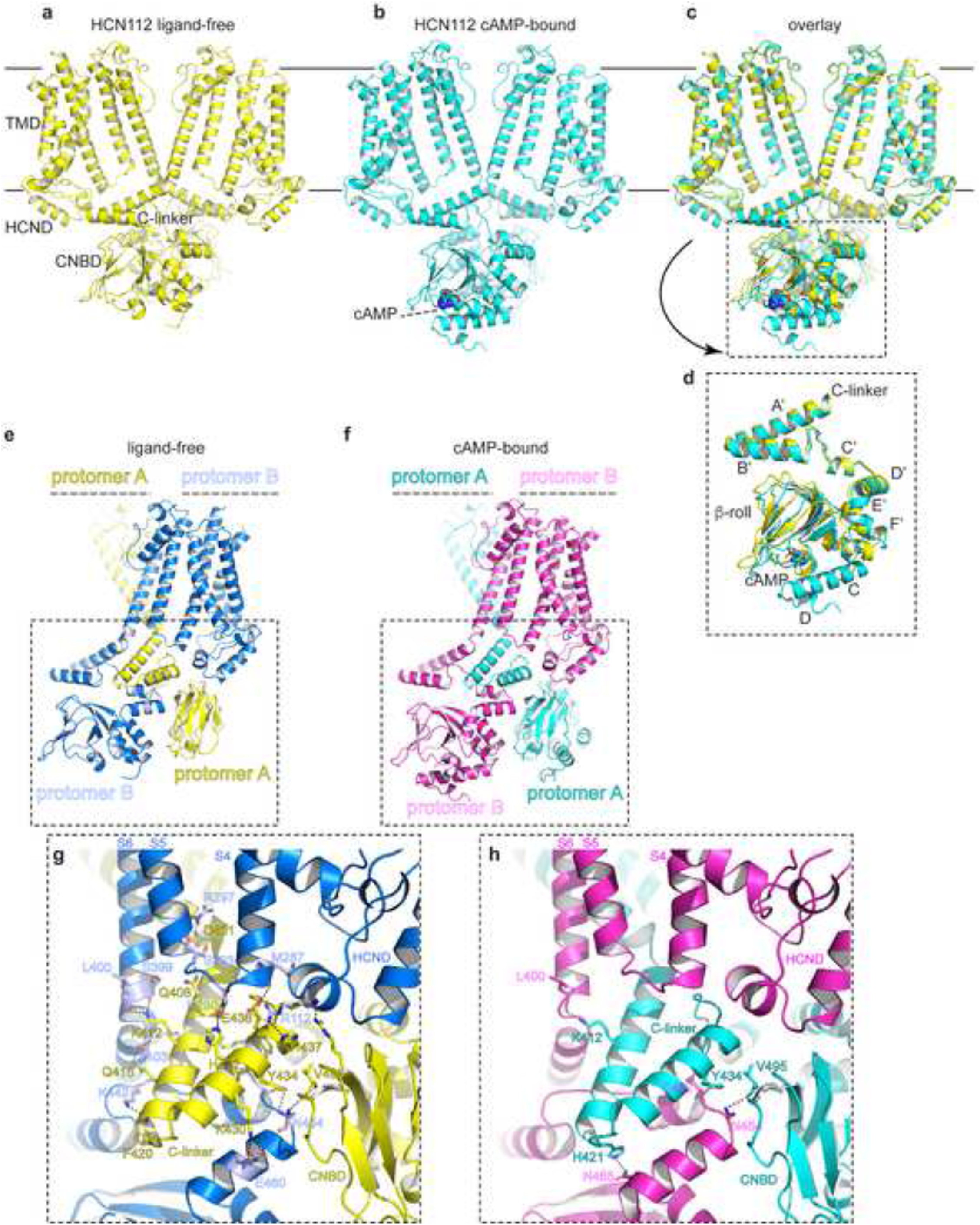
cAMP binding causes a conformational change of the C-linker and CNBD, and breaks inter-chain interactions in HCN112. a-c. Cartoon representation of HCN112 in the ligand-free conformation (**a**, yellow), cAMP-bound conformation (**b**, cyan) and overlay (**c**). Only 2 opposing subunits are shown for clarity. **d**. The inset shows a detailed view of conformational changes in the C-linker and CNBD. **e-h.** Interchain-interactions between promotor A (yellow) and promotor B (blue) in the HCN112 ligand-free conformation and promotor A (cyan) and protomer B (magenta) in the HCN112 cAMP-bound conformation. **g-h**. The insets show a detailed view of inter-chain interactions. Interacting residues are shown in ball and stick representation. Dashed lines show hydrogen bonds or salt bridges. Oxygen atoms are shown in red, nitrogen atoms are shown in blue.

As a result of this conformational change there is a large decrease in the number of inter-chain interactions under the form of hydrogen bonds and salt bridges. Specifically, there are 17 inter-chain H-bonds and salt bridges in the ligand-free form of HCN112 (see Table 1). Most of these interactions are broken in the cAMP-bound state except for 3 remaining H-bonds (K412-L400, Y434-E452, Y439-E452) and 3 H-bonds that are newly formed (H421-N465, V495-N454, R548-E575) (Figure 3F-H). For HCN112 in the ligand-free conformation we will only discuss those interactions that likely contribute to gating of the channel.

First, a key role appears to be played by R112 in the HCND domain. Three residues project interactions onto R112, namely H437, Q440 (in the C-linker) and R560 (in the CNBD). Second, there are 4 residues at the bottom of S4 and the S4-S5 linker that interact with the C-linker. These include M287 (S4), D290 and S293 (S4-S5 linker), and R297 (S5). Lastly, two residues at the bottom of S6, namely S399 and L400 interact with K412 in the C-linker.

In summary, our interpretation is that there are tight inter-chain interactions in the CNBD and C-linker, also involving the HCND domain and S4-S5 linker, and that these interactions are broken upon cAMP-binding. The breaking of these interactions could then contribute to the facilitation of channel opening.

### Deletion of the C-linker and CNBD causes partial channel opening in the HCN1ΔC channel

It has been shown that the intracellular channel region is not only responsible for mediating cAMP regulation, but also for the integrity of the channel gate, which is formed by the intracellular helix bundle of the S6 segments. Previous studies proposed a gating mechanism in which the C-linker disk plays an important role in keeping the gate closed at depolarized voltages: (1) by forming intense contacts with the unusual S4 helix^3^, and (2) by contacting the HCN domain of opposite subunits^30^. These interactions help to stabilize the gate in a closed right-handed helix bundle. For channel opening, the CL-CNBD has to perform a leftward rotation to unwind the right-handed S6 helix bundle gate. Such a leftward rotation is suggested to be induced by the movement of the voltage sensor upon a hyperpolarizing voltage jump. Cyclic nucleotide binding can cause a CL-CNBD leftward rotation, too. However, this cAMP-triggered rotation is not sufficient to open the channel but is supportive for voltage-induced rotation. In this sense, cAMP causes the CL-CNBD to adopt a conformation, which can be interpreted as relief of autoinhibition. To test this proposed mechanism, we took advantage of the HCN1ΔC construct, in which the C-terminal cytoplasmic domain was deleted^7^. The deletion spans two domains, the C-linker and the CNBD, starting from D401 till the last residue of HCN1 (Figure 1A-E). From the mechanism described above we expect that the gate and the pore move towards their open configurations when the CL-CNBD is missing. Gating of HCN channel is activated by membrane hyperpolarization and is also modulated by cAMP. It has been shown that the cyclic nucleotide–binding domain has an inhibitory effect on channel gating that is relieved by cAMP binding^7^. Such a relieving effect is reflected by an increase in current amplitude at a given voltage, by a shift of the voltage of half-maximum activation, *V*_1/2_, to more depolarized values, and by an acceleration of activation kinetics and a deceleration of deactivation kinetics^33^.

Here, we determined the cryo-EM structure of HCN1ΔC, which reached an overall resolution of 3.96Å (Figure 4). HCN1ΔC shows an RMSD value of 1.114Å and 1.096Å when aligned to the TMD domain of ligand-free HCN112 and cAMP-bound HCN112, respectively. Our HCN1ΔC structure shows a movement of pore helices S6 and S5, as shown in the inset in Figure 4 and supplementary movie S3. Overlay of cryo-EM densities of ligand-free HCN112 and HCN1ΔC (bottom left inset) clearly shows this redistribution of helices and inner sidechains along the pore.

**Figure 4.**
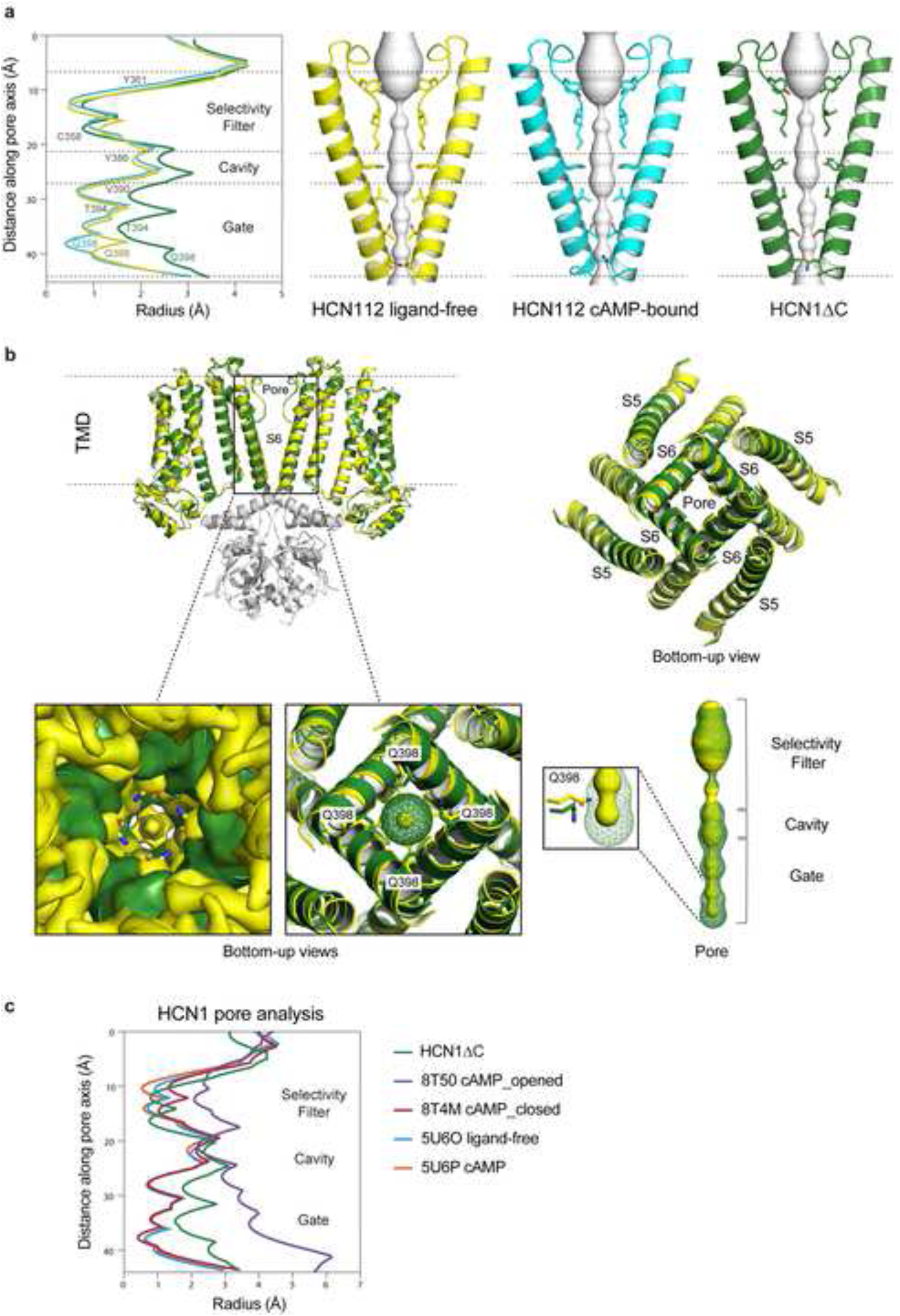
**Relief from auto-inhibition in HCN1**Δ**C reveals a pore in an intermediately open conformation. a.** Pore radius calculations for HCN112 ligand-free (yellow), HCN112 cAMP-bound (cyan), and HCN1ΔC (green) are shown on the graph (left). The corresponding structures of the pore region (A and C chains, residues 358 to 401) are shown as cartoons (right), with the sidechains of key pore-lining amino acids shown as licorice sticks. The selectivity filter, cavity, and gate regions are indicated on both the graph (left) and structures (right) with dotted lines. Grey spheres within the structures represent the calculated pore. **b.** Overlay of two subunits of HCN112 ligand-free (yellow) and HCN1ΔC (green), is shown as cartoons on the top left, with a zoomed bottom-up view on the top right. The C-linker and CNBD domain from the HCN112 ligand-free structure are colored grey. A zoomed-in view of the pore region in the overlapped structures is displayed in the bottom panels. The bottom left panel shows the overlapped structures with their corresponding cryo-EM maps, while the bottom right panel focuses on displaying the calculated pore radius within the structures (yellow sphere and green mesh) and highlighting the residue Q398 for all S6 helices. A side view of the calculated pore for both structures is included in the bottom-right corner, emphasizing the position of Q398 in the gate region. **c.** Pore radius calculations of the HCN1ΔC structure, compared to published HCN1 structures in different conformational states, are plotted. The approximate locations of the selectivity filter, cavity, and gate regions are indicated for reference. Comparison of the pore radius in the closed ligand-free HCN1 conformation (pdb code 5u6o), the closed cAMP-bound HCN1 conformation (pdb code 5u6p), the closed HCN1 F186C S264C bound to cAMP (pdb code 8t4m), the open HCN1 F186C S264C bound to cAMP (pdb code 8T50) and the intermediately open HCN1ΔC.

**Figure 5.**
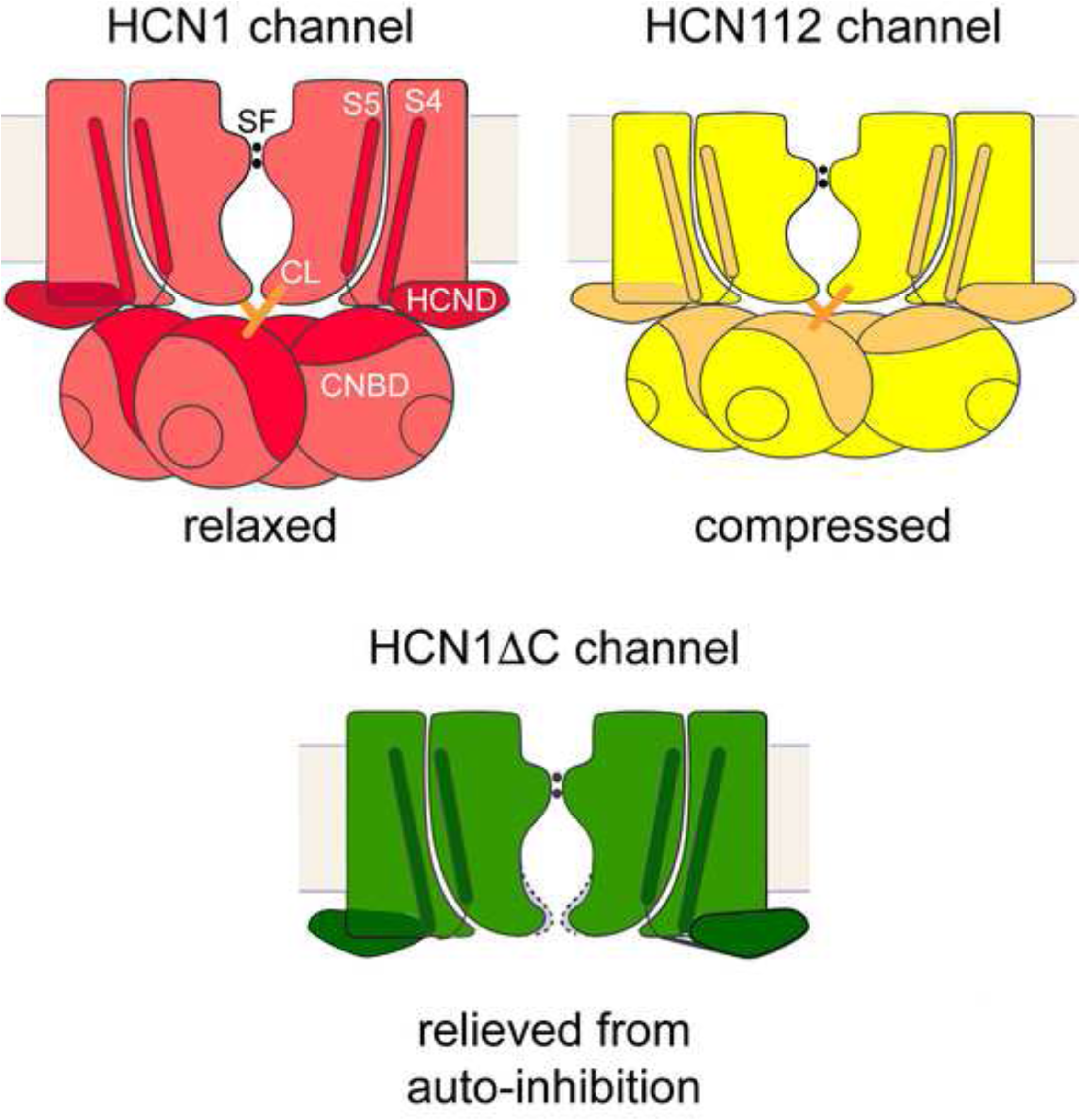
**Models of domain compression in HCN112 and relief from auto-inhibition in HCN1**Δ**C.** The models illustrate the domain compression in HCN112 (yellow) upon comparison with HCN1 in the closed ligand-free conformation (red). The green model shows relief from auto-inhibition in HCN1ΔC resulting in a partially open pore conformation. The figure was adapted from Lee and MacKinnon (2017) with permission from Elsevier.

Determinant aspects confirming the widening of the pore in HCN1ΔC are sidechains of V390, T394 and Q398. Interestingly, these residues have also been shown before to be key elements in the inner channel gate^3^. The long sidechain of Q398 at the intracellular gate of the pore tilts away from the inner pore cavity, widening the diameter of the intracellular gate of HCN1ΔC (colored in green mesh - bottom right inlet). The HCN1ΔC pore diameter is close to double in size of the HCN112 pore, which is depicted in yellow colour and overlayed in the bottom-up view in Figure 4A and B. These small differences in conformation of S6-S5 helices in HCN1ΔC led the overall pore diameter to be halfway toward being fully open (∼3Å vs ∼5Å) and thus resembles the open HCN1 conformation as well as the rabbit HCN4 open pore conformation (Figure 4B-C).

In summary, we hypothesized that the S6-S5 and S4 helices movements cause a loss of contacts between these pore helices and the C-linker, as well as between the HCND and the CNBD. We therefore suggest that these rearrangements reflect the transition state towards an open pore in HCN1, triggering a release of constraints and allowing the pore to open. We also propose that our HCN1ΔC structure, with its deletion reflecting an absence of auto-inhibition, mimics a transitioning-hyperpolarized state, that most likely precedes the hyperpolarized state described in Lee and MacKinnon^4^.

## MATERIALS AND METHODS

### Constructs design

All HCN1 constructs were engineered in the background of the human HCN1_EM_ plasmid made by Lee and MacKinnon^3^ and was cloned into a pEG-BacMam vector^34^ for expression in mammalian cells and which tags the protein of interest with both N-His10, eGFP and 3C protease site. The following constructs were generated on top of the HCN1_EM_ constructs: 1. Chimera HCN112 contains region Met1-Leu400 from human HCN1 and Asp401-Leu658 from human HCN2, with the region between Ala705-Ser866 of HCN2 which was deleted, while the HCN2 extreme C-terminal region including the SNL tripeptide sequence was preserved Ala867-Leu889 (residue numbering according to full-length human HCN2); 2. HCN1ΔC corresponds to the HCN1 gene covering the sequence between Met1-Asp401, with a deletion that spans the entire C-terminal region.

### Electrophysiological recordings

For expression in *Xenopus* oocytes HCN112 and HCN1ΔC were subcloned into pGEM-HE using *Xba*I and *Hind*III restriction sites. Plasmids were linearized with *Nhe*I and transcribed with T7 RNA polymerase using the mMessage mMachine T7 transcription kit. The RNA quality was verified by loading RNA samples on 1% agarose gel.

Oocytes were surgically removed from adult female South African claw frogs *Xenopus laevis* under anesthesia with 0.3% tricaine methanesulfonate (MS-222) (Pharmaq Ltd. Fordingbridge, United Kingdom). After removal, the oocytes were treated with collagenase A (3 mg/ml; Roche, Grenzach-Wyhlen, Germany) for 105 min in Ca^2+^-free Barth’s solution containing (in mM) 82.5 NaCl, 2 KCl, 1 MgCl_2_, and 5 Hepes, pH 7.5. Oocytes of stages IV and V were manually dissected and injected with cRNA encoding either HCN112 or HCN1ΔC channels. After injection with cRNA, the oocytes were incubated at 18°C for 2–6 days in Barth’s solution containing (in mM) 84 NaCl, 1 KCl, 2.4 NaHCO_3_, 0.82 MgSO4, 0.41 CaCl_2_, 0.33 Ca(NO3)_2_, 7.5 TRIS, pH 7.4. Oocytes harvested in our lab were complemented with ready-to-use oocytes purchased from Ecocyte Bioscience (Dortmund, Germany). The surgery procedures were carried out in accordance with the German Animal Welfare Act with the approval of the Thuringian State Office for Consumer Protection on 30.08.2013 and 09.05.2018.

Macroscopic currents were recorded using the inside-out configuration of the patch-clamp technique. All measurements were started after a delay of 3.5 min to minimize run-down phenomena. Patch pipettes were pulled from quartz tubings with outer and inner diameters of 1.0 and 0.7 mm (VITROCOM, New Jersey, United States), respectively, using a laser puller (P-2000, Sutter Instrument, Novato, United States). The pipette resistance was 1.2–2.1 MOhm. The bath solution contained (in mM) 100 KCl, 10 EGTA, and 10 Hepes, pH 7.2, and the pipette solution contained (in mM) 120 KCl, 10 Hepes, and 1.0 CaCl_2_, pH 7.2. For parts of the experiments, a saturating concentration of 10 µM cAMP (BIOLOG LSI GmbH & Co. KG, Bremen, Germany) was applied with the bath solution. A HEKA EPC 10 USB amplifier (Harvard Apparatus, Holliston, United States) was used for current recording. Pulsing and data recording were controlled by the Patchmaster software (Harvard Apparatus, Holliston, United States). The sampling rate was 5 kHz. The holding potential was generally −30 mV. Maximally two membrane patches were excised from one individual oocyte.

### Protein expression, purification and sample preparation for Cryo-EM

The different HCN constructs were expressed in HEK293S GnTI^-^ cells (Goehring et al., 2014) at 37°C and to a density of ∼3x10^6^ cells/ml for ∼72h, after addition of baculoviruses carrying HCN1 genes (10% v/v). After 15h, 10 mM sodium butyrate was added to cells and the culture temperature was shifted to 30°C for ∼72 h, after which cells were harvested and pellets were flash frozen and stored at −80°C.

Cell pellets were resuspended in 30% glycerol for 10 min and lysed by gentle homogenization in 10 mM Tris pH 8.0, 20 mM KCl, 2 mM DTT, 0.1 mg/ml DNAse, 0.5 mM MgCl_2_, 2 μg/ml leupeptin, 10 μg/ml aprotinin, 1 μg/ml pepstatin, 50 μg/ml benzamidine HCl, 1 mM AEBSF, 30% glycerol for 25 min at 4°C. The lysate was spun at 39,800g for 35 min at 4°C to sediment crude membranes. The membrane pellet was homogenized and solubilized in the extraction buffer (20 mM Tris pH 8.0, 300 mM KCl, 2 mM DTT, 2 μg/ml leupetin, 10 μg/ml aprotinin, 1 μg/ml pepstatin, 50 μg/ml benzamidine HCl, 1 mM AEBSF, 10 mM LMNG + 2 mM CHS) for 2 h. Solubilized membranes were clarified by centrifugation at 39,800g for 35 min, 4°C. The supernatant was incubated with the GFP nanobody-coupled sepharose resin prepared in-house^35^ for 1.5 h at 4°C and subsequently washed for 10 column volumes with wash buffer (20 mM Tris pH 8.0, 300 mM KCl, 2 mM DTT, 2 μg/ml leupetin, 10 μg/ml aprotinin, 1 μg/ml pepstatin, 50 μg/ml benzamidine HCl, 1 mM AEBSF, 0.05% digitonin). An extra 5 column volumes of buffer with 1 mM EDTA was used to incubate the proteins of interest overnight at 4°C with PreScission protease to cleave the GFP tag. HCN variants were eluted straight on a Glutathione Sepharose 4B column and washed out with wash buffer. The eluted proteins were concentrated and purified further through a Superose 6 10/300 column using gel filtration buffer (20 mM Tris pH 8.0, 150 mM KCl, 2 mM DTT, 2 μg/ml leupeptin, 10 μg/ml aprotinin, 1 μg/ml pepstatin, 50 μg/ml benzamidine HCl, 1 mM AEBSF, 0.05% digitonin). Peak fractions of interest were pooled, concentrated to 3-4 mg/ml, flash frozen in liquid nitrogen and stored at −80C.

Once thawed, the purified proteins were spun at 13,000 rpm for 5 min at 4°C and the supernatant was used for grid preparation. 0.4 mM cAMP was added to 3.5 mg/ml HCN112 in order to perform single particle analysis on the HCN112 cAMP-bound sample. 3 μl of the purified HCN variants were applied on glow discharged AuFoil grid R2/2 200 mesh within a Vitrobot IV (FEI) operated a 4°C, 100% humidity (blot time 3 s, waiting time 3 s) and plunged into liquid ethane.

### Cryo-EM data acquisition

Two data acquisitions were acquired using two Titan Krios transmission electron microscopes (FEI) operated at 300keV with energy filter, equipped with Falcon 4 electron detector (ABSL, Leeds, UK) at a nominal magnification of 130k, with a pixel size of 0.91Å (HCN112+cAMP and HCN1ΔC). Two other datasets, later merged as one single dataset, were acquired at a nominal magnification of 96k, with a pixel size of 0.81Å (ligand-free HCN112).

The nominal defocus range chosen was between the following values and changing in increments of 0.2: for apoHCN112 this was 1.6 - to 3.2 μm, for HCN112+cAMP this was 1.2 to 3 μm, for HCN1ΔC this was 1.6 to 3.4 μm. Respectively, the total doses were: 41.42 e^-^/Å^2^ (merged ligand-free HCN112), 32.44 e^-^/Å^2^ (HCN112+cAMP) and 39.81 e^-^/Å^2^ (HCN1ΔC). The exposure time for each image was 6.26 s (ligand-free HCN112), 3.98s (HCN112+cAMP), 6.49s (HCN1ΔC), fractionated over 54, 41, 40 EER fractions, respectively. Acquisition parameters are summarized in Table S1.

### Image processing and single-particle analysis

Movies were corrected for beam-induced motion using MotionCor (relion3.1.1)^36^ and contrast transfer function parameters were estimated from motion-corrected images using CTFFIND 4.1. For all datasets, particle picking was performed in crYOLO v1.61_GPU^37^ using an integrated general model and cutoff 0.1 and JANNI denoised. Particles were extracted in Relion with a box size of 350px. The particle.star file was then imported in cryoSPARC v.3.2.0^38^, where 2D classification, ab-initio reconstruction and refinements were performed (Suppl Fig. 2-7 for Cryo-EM flowchart). During refinement, C4 symmetry was imposed in all datasets. The same single-particle processing was applied for all datasets, leading to a local resolution of 2.9 Å ligand-free HCN112, 2.7Å HCN112+cAMP, 3.45Å HCN1ΔC.

### Model building, refinement and validation

Manual building and real-space refinement was performed using HCN1 (PDB 5u6o) as starting model for the transmembrane region (TM) and HCN2 crystal structure (PDB 3u10) for the cytoplasmic domain (CD) in Coot v0.8.9.2^39^ and then in Phenix v1.19.2^40^. These homology models were docked into the density map of the ligand-free HCN112 using UCSF chimera^41^ in order to build a starting model for one protomer of the chimeric HCN112. A tetramer model of the channel was obtained by fitting copies of the same model in the C4 symmetric map of the tetramer. The starting model for the cAMP-bound HCN112 was the refined ligand-free HCN112. The placement of the cAMP ligand in the holoHCN112 was based on the crystal structure of the HCN2 in complex with cAMP (PDB 3u10). As a model for HCN1ΔC TM region, the cAMP-bound HCN112 was used but manual rebuilding was necessary for amino acid fragments that undergo conformational changes i.e. S6, S5 and HCN domain.

The final ligand-free structure includes residues 97–240, and 252–579, whereas the cAMP-bound structure includes additional residues 580–605 and part of the E-helices built as a poly-Ala helix. For the HCN1ΔC structure building was more challenging due to the lower quality of the cryo-EM map, but residues of the pore region (361-398) could be built using the HCN112+cAMP structure as a reference thus allowing analysis of the pore radius profile. The quality of the final models was evaluated by MolProbity^42^ and EMRinger^43^ (Table S1). Figures were prepared using Chimera^41^, ChimeraX^44^ and PyMOL Molecular Graphics System (v1.8, Schrodinger, LLC). Pore analysis was performed in HOLE v2.2.005^45^ after superposition of all protein structures of interest. Interactions between residues were calculated using Contact/Act in the CCP4 suite^46^ using a maximum distance cut-off of 3.8 Å. H-bonds and salt bridges were considered between 2.6-3.4 Å and van der Waals interactions were considered between 3.2-3.8 Å.

## DATA RESOURCES

Cryo-EM density maps of HCN112 apo, HCN112+cAMP, and HCN1ΔC have been deposited in the electron microscopy data bank under accession code EMD-53516, EMD-53514 and EMD-53515, respectively. Atomic coordinates of HCN112 apo, HCN112+cAMP and HCN1ΔC have been deposited in the protein data bank under accession code 9R1V, 9R1T and 9R1U, respectively.

## CONTRIBUTIONS

VZ performed protein expression, purification, sample and grid preparation, collected and processed Cryo-EM data. MB performed cloning and helped with purification. RG helped with grid preparation, collection and processing of cryo-EM data. VZ, JMKF, SDG, RGE and CU analyzed the data and prepared figures. CU designed and guided the project. VZ and CU wrote the manuscript with input from other authors. All authors have reviewed the manuscript.

## ACKNOWLEDGEMENTS

The research by CU was supported by KU Leuven grant C14/23/128 and grants G0C1319N, G087921N from FWO-Vlaanderen. We thank Dr. Rebecca Thompson and all the Cryo-EM team at Astbury Center for Molecular and Structural Biology, Leeds, UK for access to and technical support for Krios Titans. The access was granted by iNext-Discovery. VZ was funded by Fund for Scientific Research Flanders Postdoctoral Fellowship (FWO 12W4618N).

## CELL PRESS DECLARATION OF INTERESTS POLICY

Transparency is essential for a reader’s trust in the scientific process and for the credibility of published articles. At Cell Press, we feel that disclosure of competing interests is a critical aspect of transparency. Therefore, we require a “declaration of interests” section in which all authors disclose any financial or other interests related to the submitted work that (1) could affect or have the perception of affecting the author’s objectivity or (2) could influence or have the perception of influencing the content of the article.

### What types of articles does this apply to?

We require that you disclose competing interests for all submitted content by completing and submitting the form below. We also require that you include a “declaration of interests” section in the text of all articles even if there are no interests to declare.

### What should I disclose?

We require that you and all authors disclose any personal financial interests (e.g., stocks or shares in companies with interests related to the submitted work or consulting fees from companies that could have interests related to the work), professional affiliations, advisory positions, board memberships (including membership on a journal’s advisory board when publishing in that journal), or patent applications and/or registrations that are related to the subject matter of the contribution. As a guideline, you need to declare an interest for (1) any affiliation associated with a payment or financial benefit exceeding $10,000 p.a. or 5% ownership of a company or (2) research funding by a company with related interests. You do not need to disclose diversified mutual funds, 401ks, or investment trusts.

Authors should also disclose relevant financial interests of immediate family members. Cell Press uses the Public Health Service definition of “immediate family member,” which includes spouse and dependent children.

### Where do I declare competing interests?

Competing interests should be disclosed on this form as well as in a “declaration of interests” section in the manuscript. This section should include financial or other competing interests as well as affiliations that are not included in the author list. Examples of “declaration of interests” language include:

“AUTHOR is an employee and shareholder of COMPANY.”

“AUTHOR is a founder of COMPANY and a member of its scientific advisory board.”

*NOTE*: Primary affiliations should be included with the author list and do not need to be included in the “declaration of interests” section. Funding sources should be included in the “acknowledgments” section and also do not need to be included in the “declaration of interests” section. (A small number of front-matter article types do not include an “acknowledgments” section. For these articles, reporting of funding sources is not required.)

### What if there are no competing interests to declare?

If you have no competing interests to declare, please note that in the “declaration of interests” section with the following wording:

“The authors declare no competing interests.”

---

**Supplementary Fig. 1.**
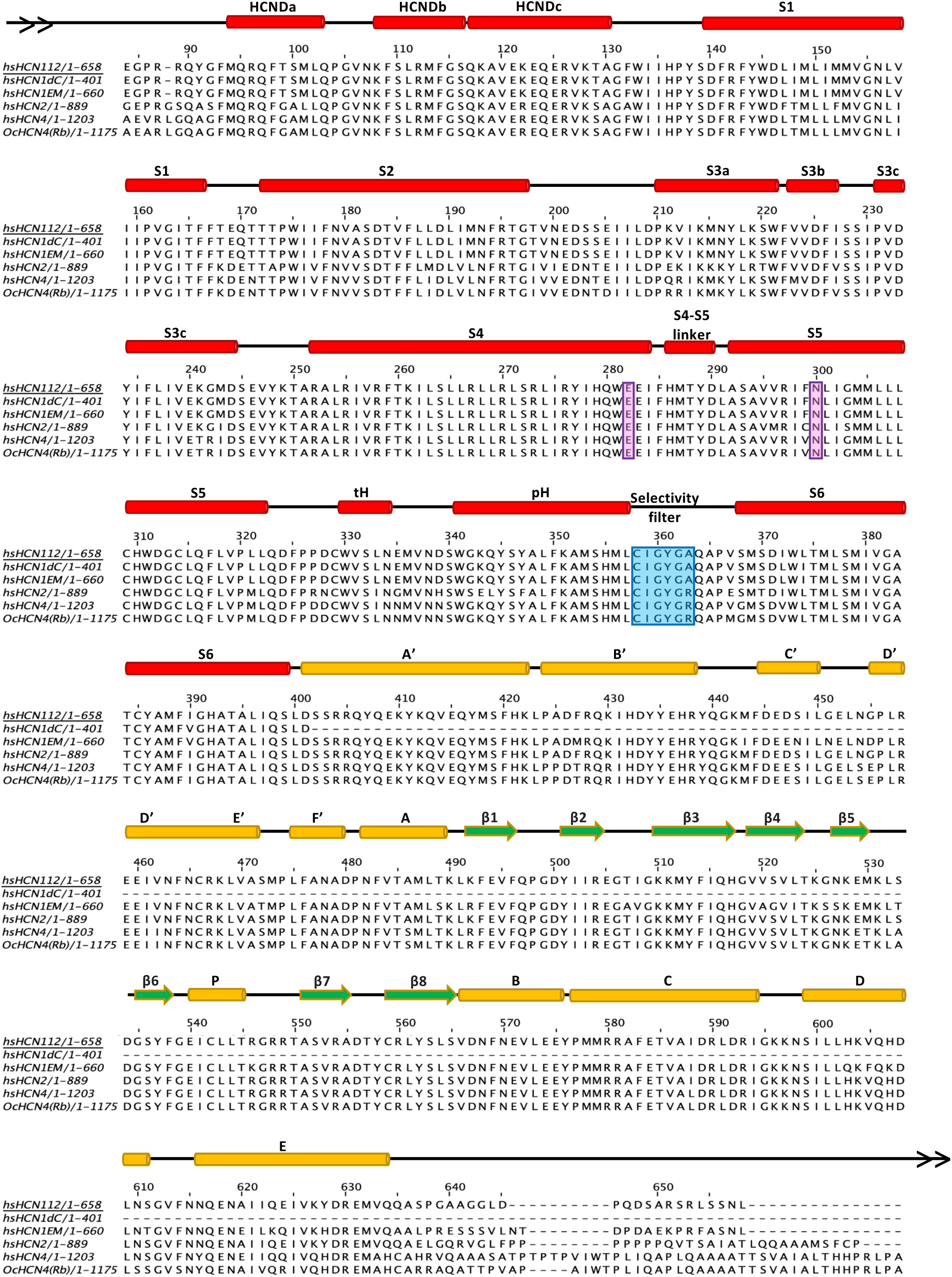
Sequence alignment and secondary structure elements of HCN channel variants. Human sequence of HCN1 EM construct, HCN112 and HCN1ΔC are shown and aligned between them and to human HCN2 (Uniprot code: Q9UL51) and HCN4 sequences (Uniprot code: Q9Y3Q4). In addition, rabbit HCN4 sequence is also represented and aligned (Uniprot code: Q9TV66). Secondary structure elements are denoted by cylinders (-helices, red), arrows (-strand, green) and lines (loops, black). Selectivity filter residues are highlighted with a cyan box. Residues that are forming a polar contact between S4 (E403) and S5 (N421) in the closed pore conformation are highlighted with a magenta box. This polar contact is absent in the open pore conformation in HCN1ΔC. N- and C-terminus have been omitted for all the sequences shown.

**Supplementary Fig. 2.**
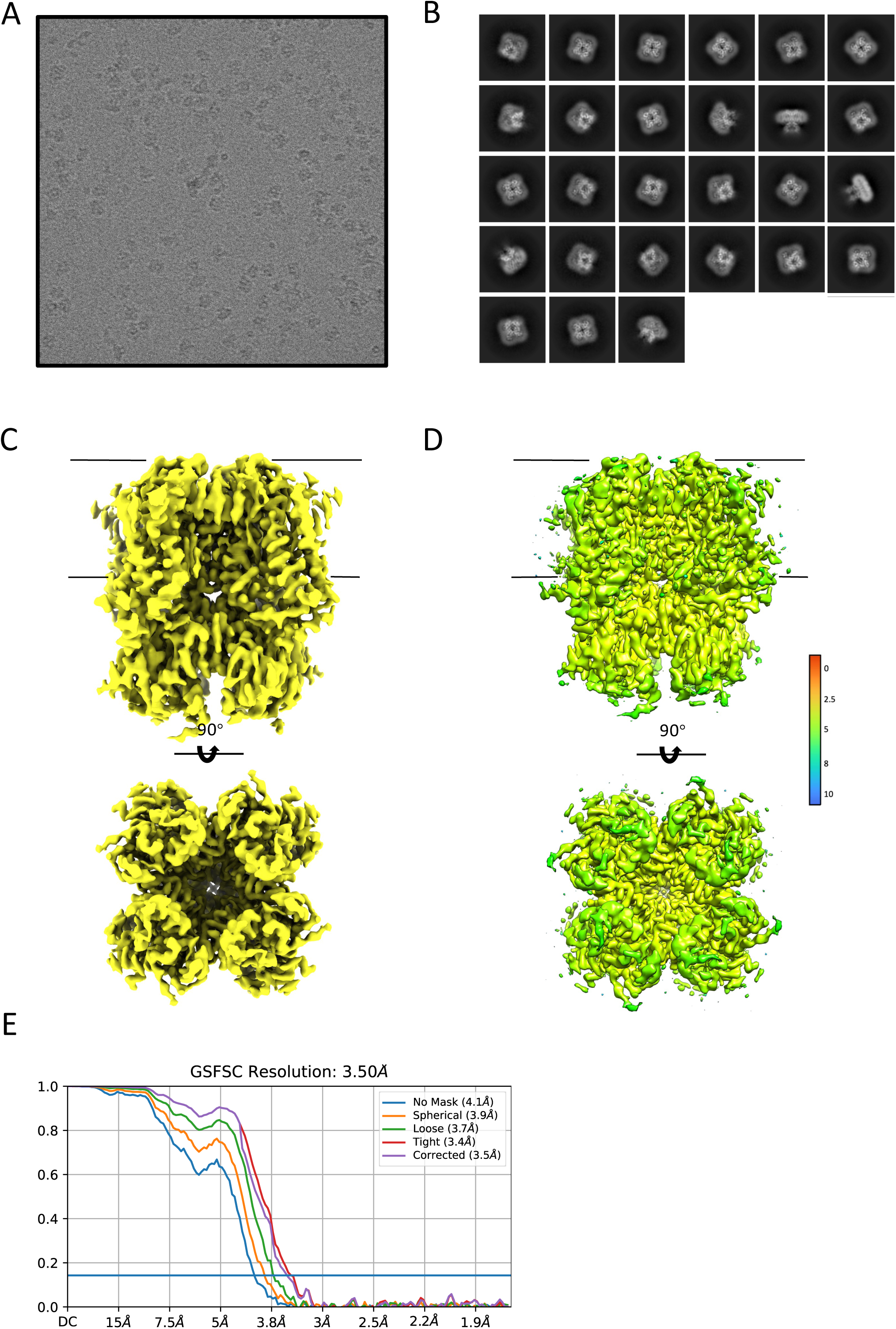
Single-Particle Cryo-EM of HCN112, ligand-free state. A) Representative raw micrograph of the HCN112 in its ligand-free state. B) Selected 2D class averages. C) HCN112 density map sharpened with a *B*-factor of −177.4 Å^2^. D) Local resolution of the density map estimated using Cryosparc_v.0.3 varying between 3.53-2.9 Å and colored with Chimera software. E) Fourier shell correlation (FSC) plots of HCN112 in its ligand-free structure using various masks.

**Supplementary Fig. 3.**
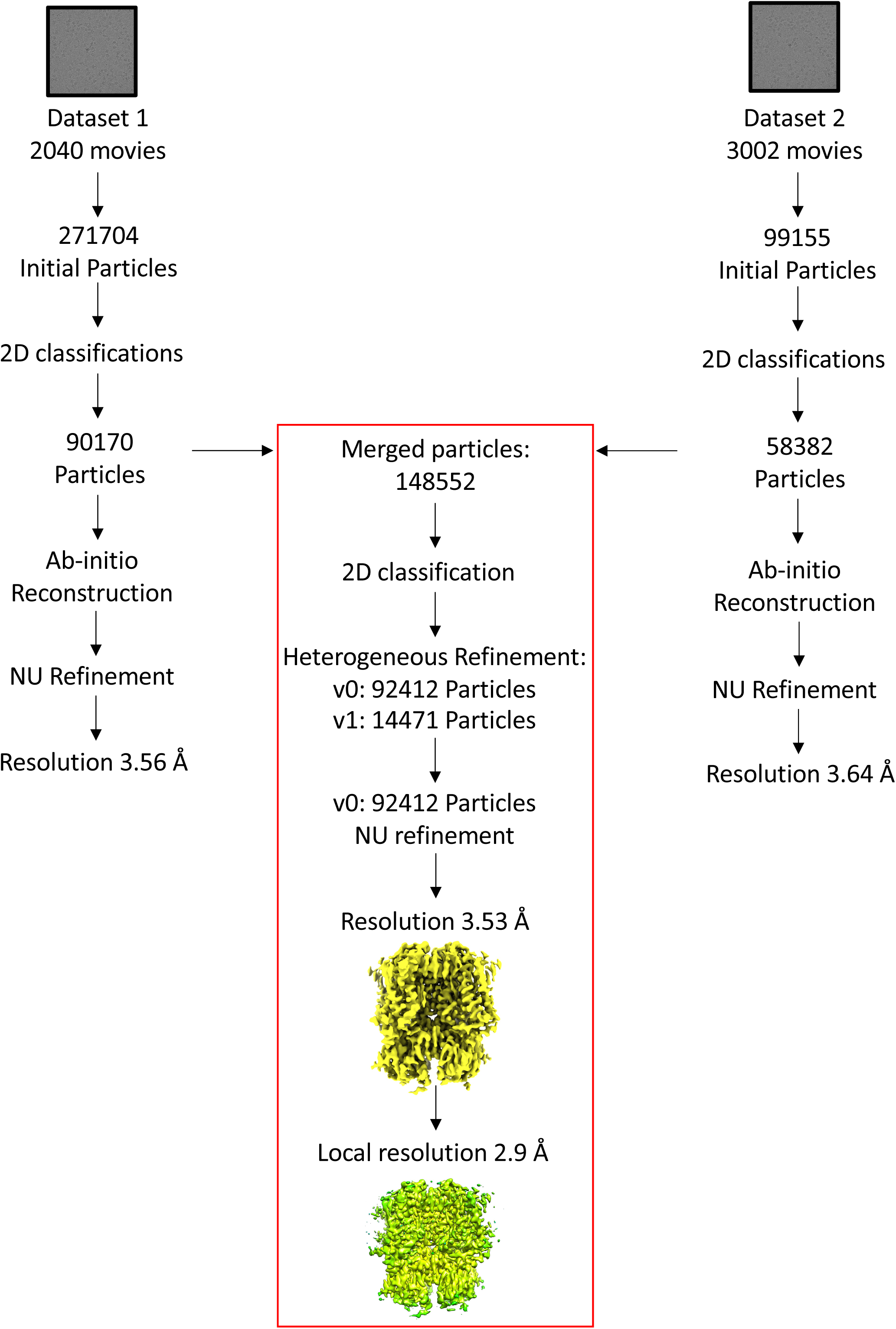
Single-Particle cryo-EM processing workflow of HCN112, ligand-free state. Processing summary for HCN112 ligand-free state. Initial processing steps that include initial number of particles used and multiple rounds of 2D classification were done in parallel, independently, on the two datasets, as well as the following steps in processing as ab-initio and refinement steps. However, the 2 datasets were merged and duplicated particles were removed before extra steps of 2D classification. NU refinement was performed only on one volume as output of a heterogeneous refinement. A resolution of 3.53 Å was achieved with 92412 particles. Local resolution of density map shown reached 2.9 Å.

**Supplementary Fig. 4.**
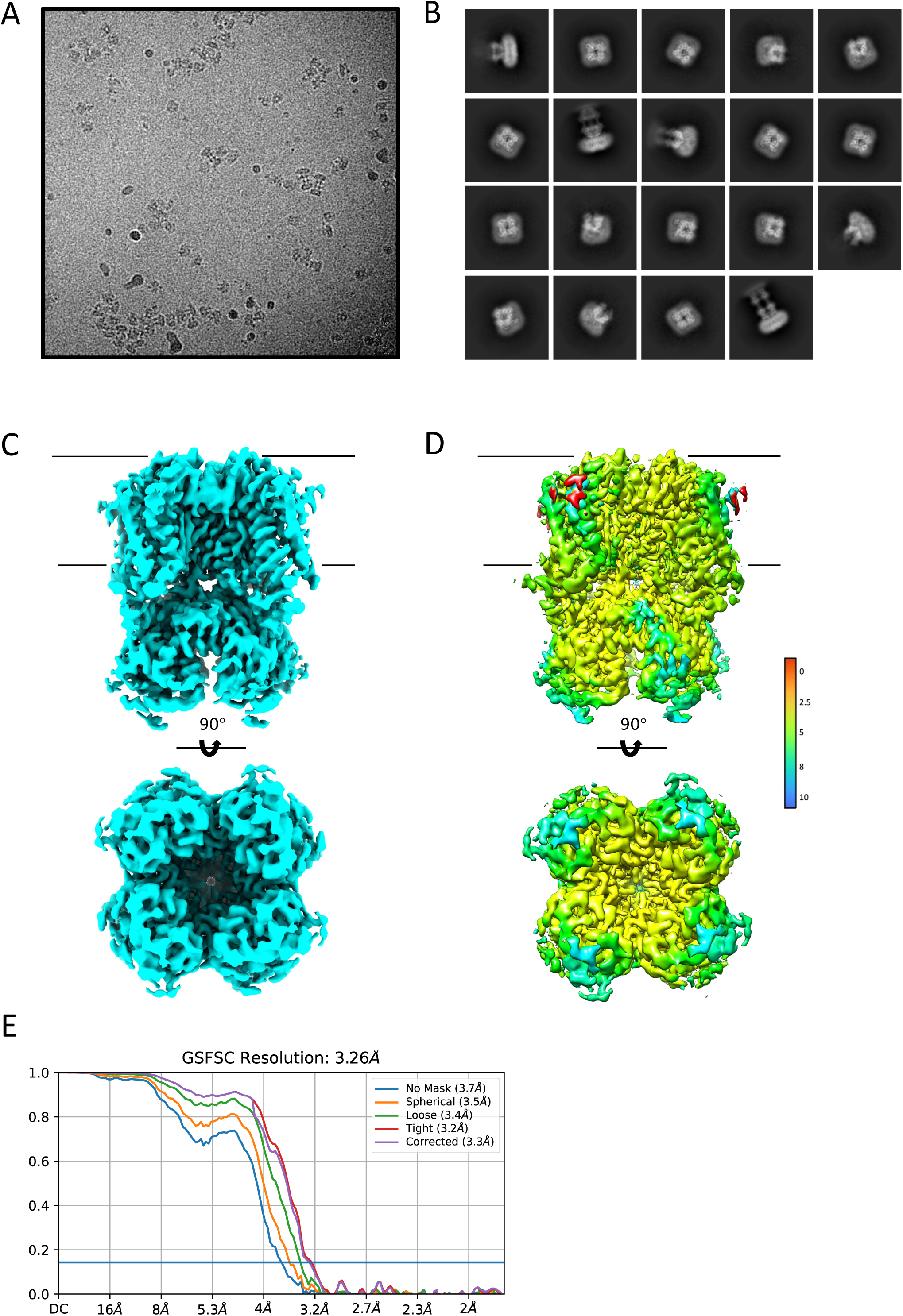
Single-Particle Cryo-EM of HCN112, cAMP-bound state. A) Representative raw micrograph of the HCN112 in its ligand-bound state with cAMP. B) Selected 2D class averages. C) HCN112 cAMP-bound density map sharpened with a *B*-factor of −147.7 Å^2^. D) Local resolution of density map estimated using Cryosparc_v.0.3 varying between 3.26-2.7 Å and colored with Chimera software. E) Fourier shell correlation (FSC) plots of HCN112 cAMP-bound structure using various masks.

**Supplementary Fig. 5.**
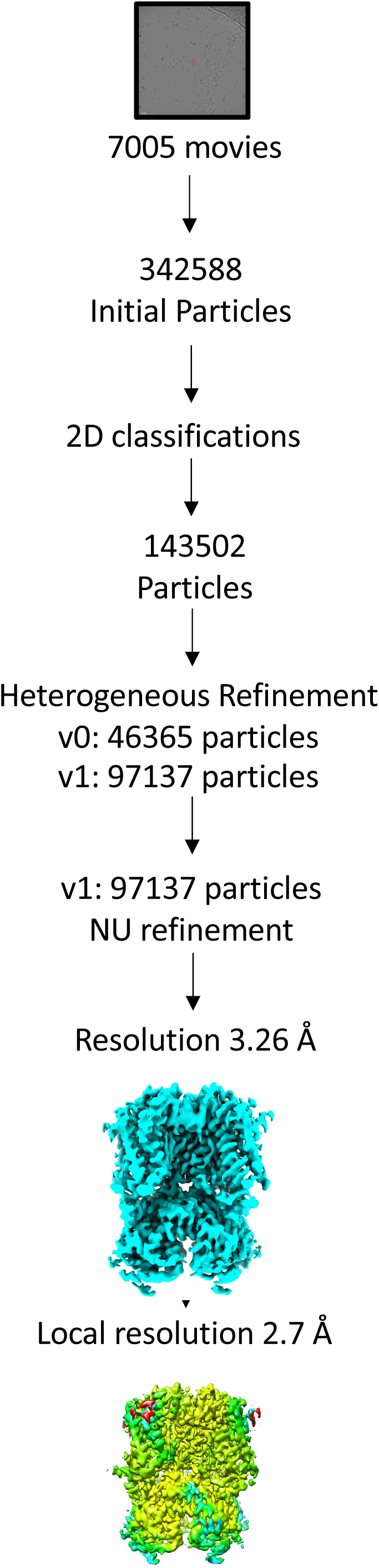
Single-Particle cryo-EM processing workflow of HCN112, cAMP-bound state. Processing summary for HCN112 cAMP-bound state. After multiple rounds of cleaning by 2D classification the dataset was further processed with 97137 particles as one volume from a heterogeneous refinement step. After NU refinement a resolution of 3.26 Å was achieved, with local resolution of 2.7 Å.

**Supplementary Fig. 6.**
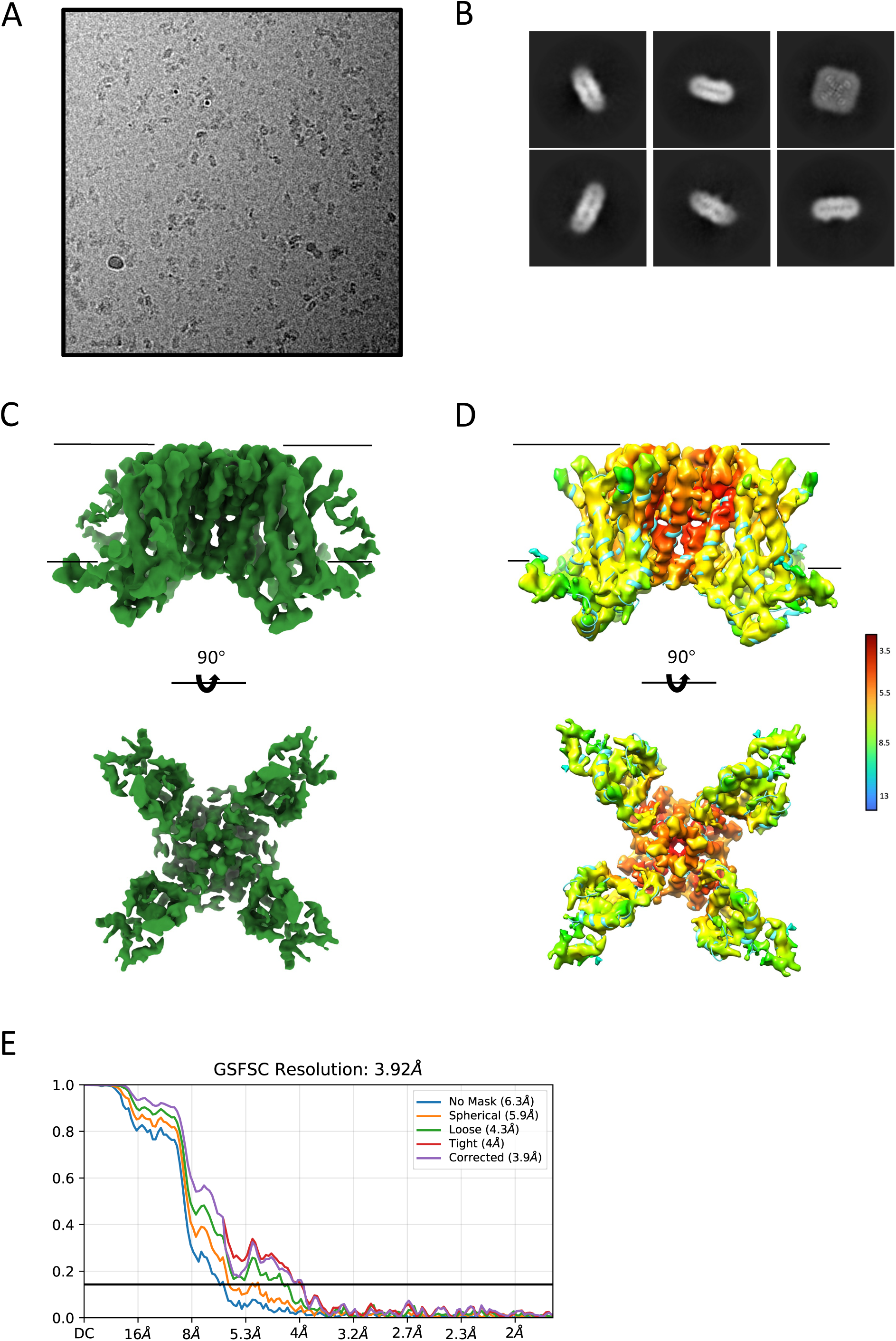
Single-Particle Cryo-EM of HCN1ΔC. A) Representative raw micrograph of the HCN1ΔC. B) Selected 2D class averages. C) HCN1ΔC density map sharpened with a *B*-factor of −140.8 Å^2^. D) Local resolution of density map estimated using Cryosparc_v.0.3 varying between 3.92-3.45 Å and colored with Chimera software. E) Fourier shell correlation (FSC) plots of HCN1ΔC structure using various masks.

**Supplementary Fig. 7.**
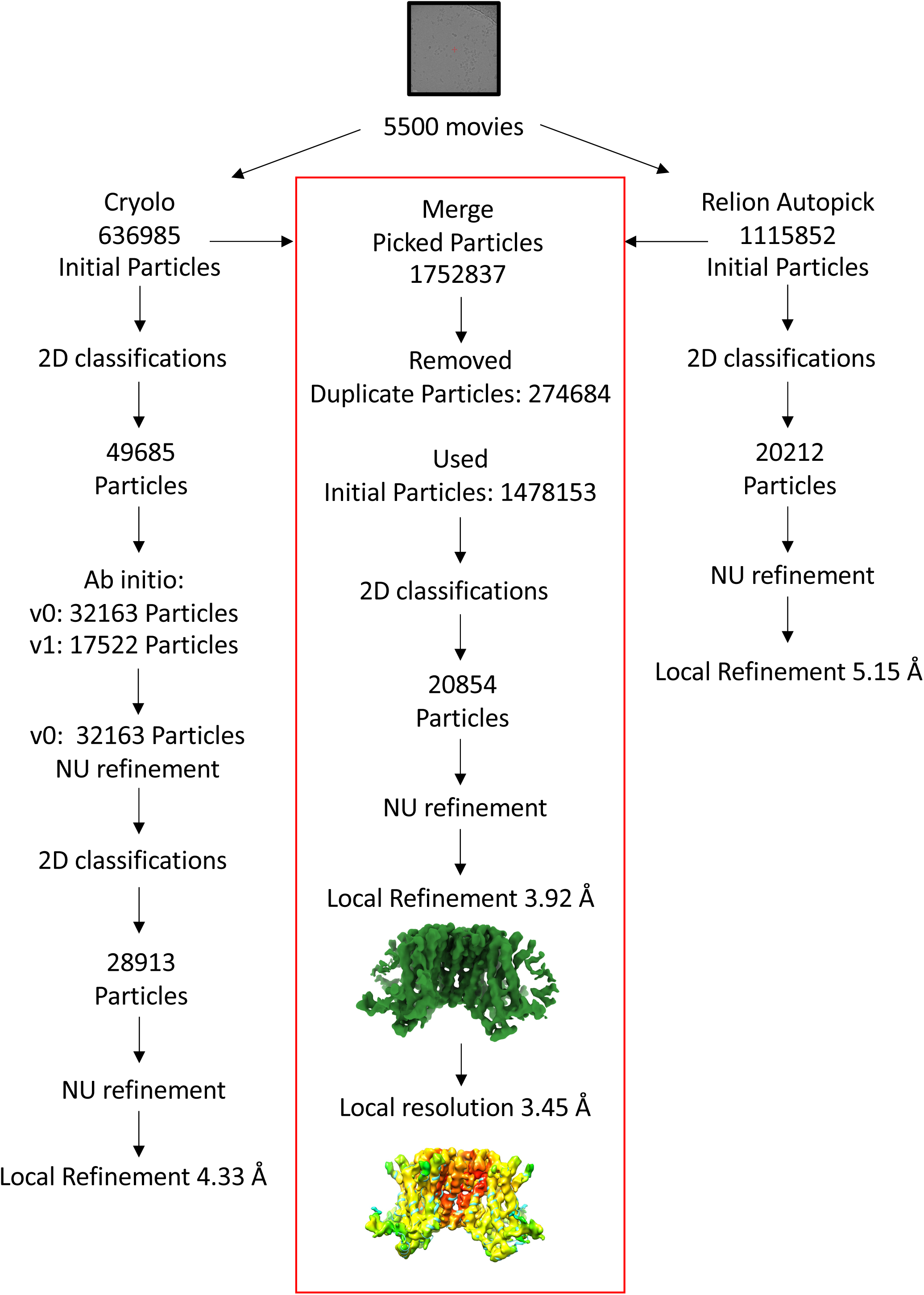
Single-Particle cryo-EM processing workflow of HCN1ΔC. Processing summary for HCN1ΔC. Initial processing steps that include picking and multiple rounds of 2D classification were done in parallel, on the unique dataset of 5500 movies, using crYOLO and Relion Autopick. On the two parallel processing, HCN1ΔC could not be resolved to high resolution, therefore picked particles were merged and duplicated removed, leading to 1478153 particles. Subsequent steps of 2D classification and refinement lead to resolution of 3.92 Å with local resolution of density map shown that reached 3.45 Å.

**Supplementary Fig. 8.**
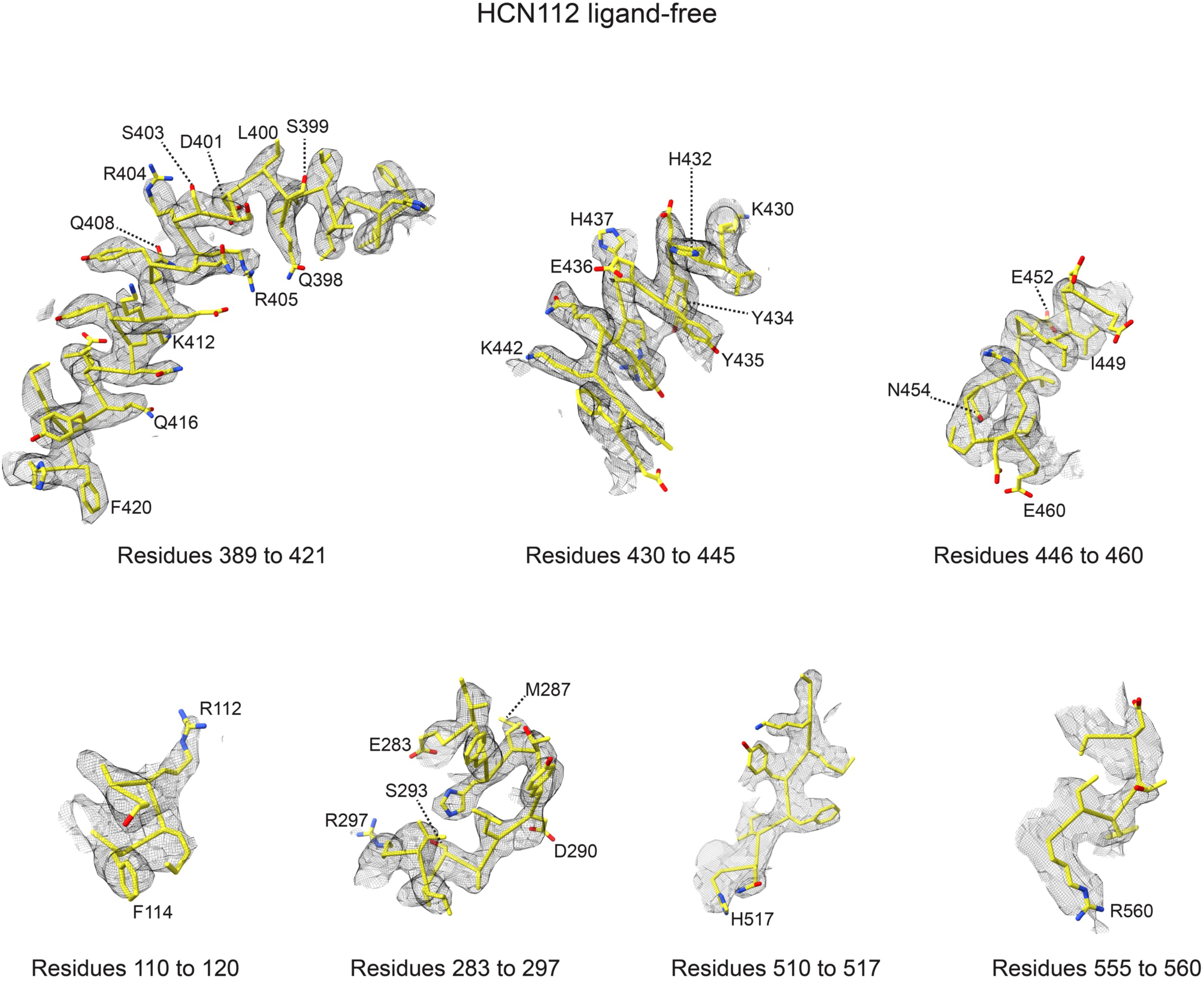
Cryo-EM density for specific residues of HCN112 ligand-free. Residues from one example subunit (chain A) are shown with their backbones and side chains (in yellow), together with their density (in grey mesh). Important residues discussed on Figure 2 are labelled. The images were generated in ChimeraX-1.8.

**Supplementary Fig. 9.**
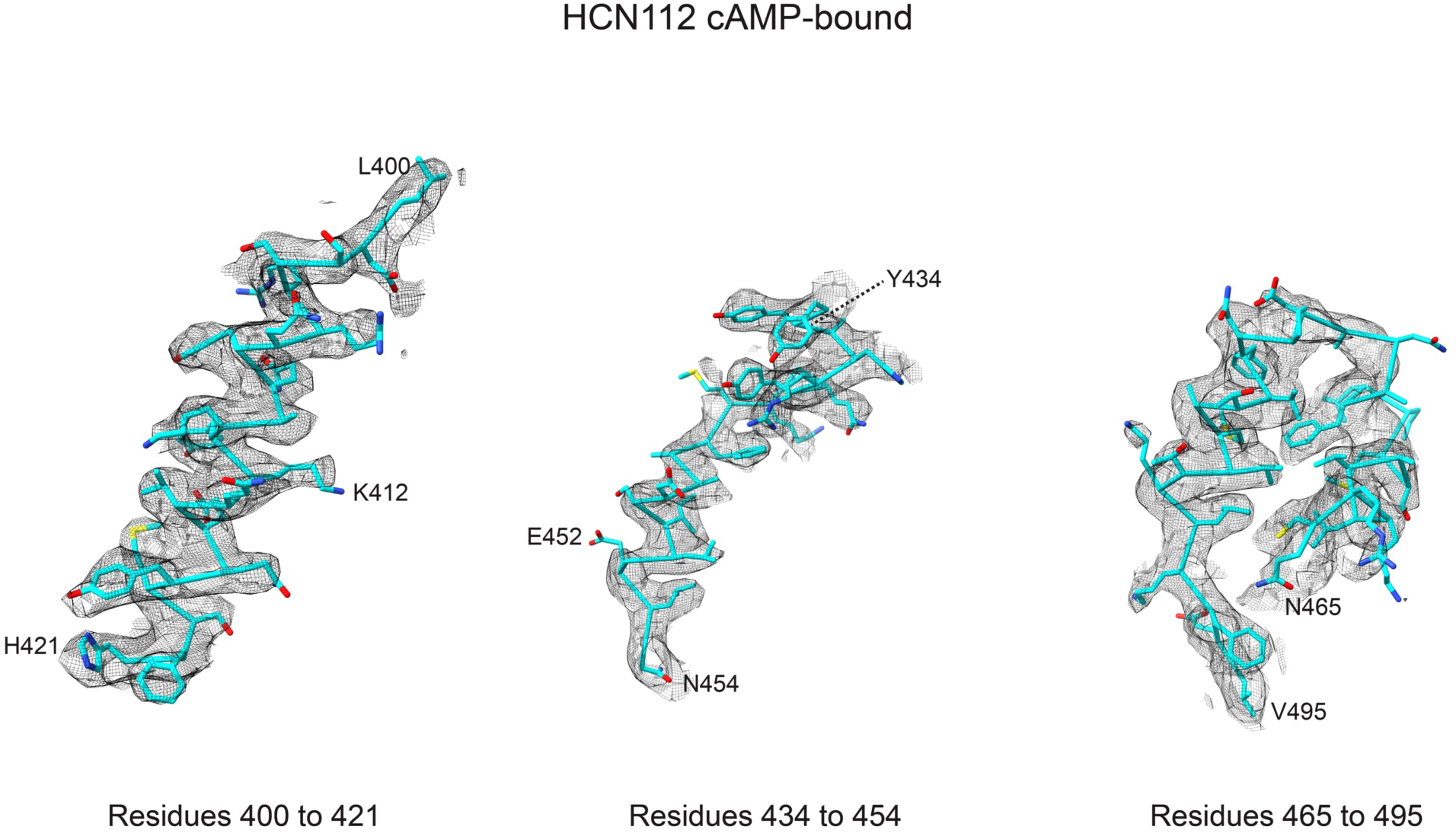
Cryo-EM density for specific residues of HCN112 cAMP-bound. Residues from one example subunit (chain A) are shown with their backbones and side chains (in cyan), together with their density (in grey mesh). Important residues discussed on Figure 3 are labelled. The images were generated in ChimeraX-1.8.

**Supplementary Fig. 10.**
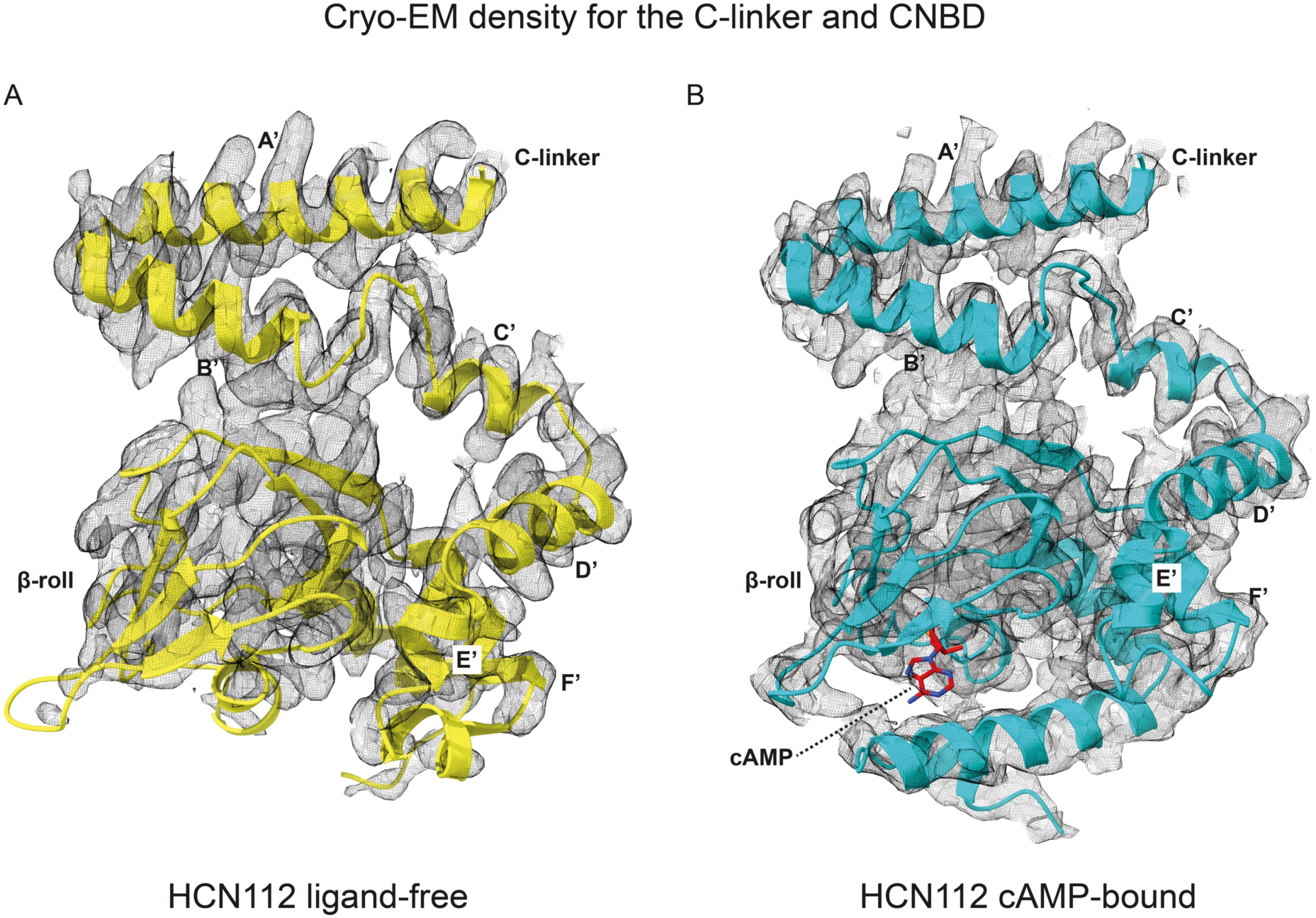
Cryo-EM density for the C-linker and CNBD domains. A) HCN112 ligand-free (yellow), and B) HCN112 cAMP-bound (cyan). The C-linker and CNBD domains from subunit A are shown as cartoon for both structures, together with the density (in grey mesh). cAMP molecule is shown as atoms and colored red in figure B. The C-linker, helices and β-roll are labelled.

**Supplementary Fig. 11.**
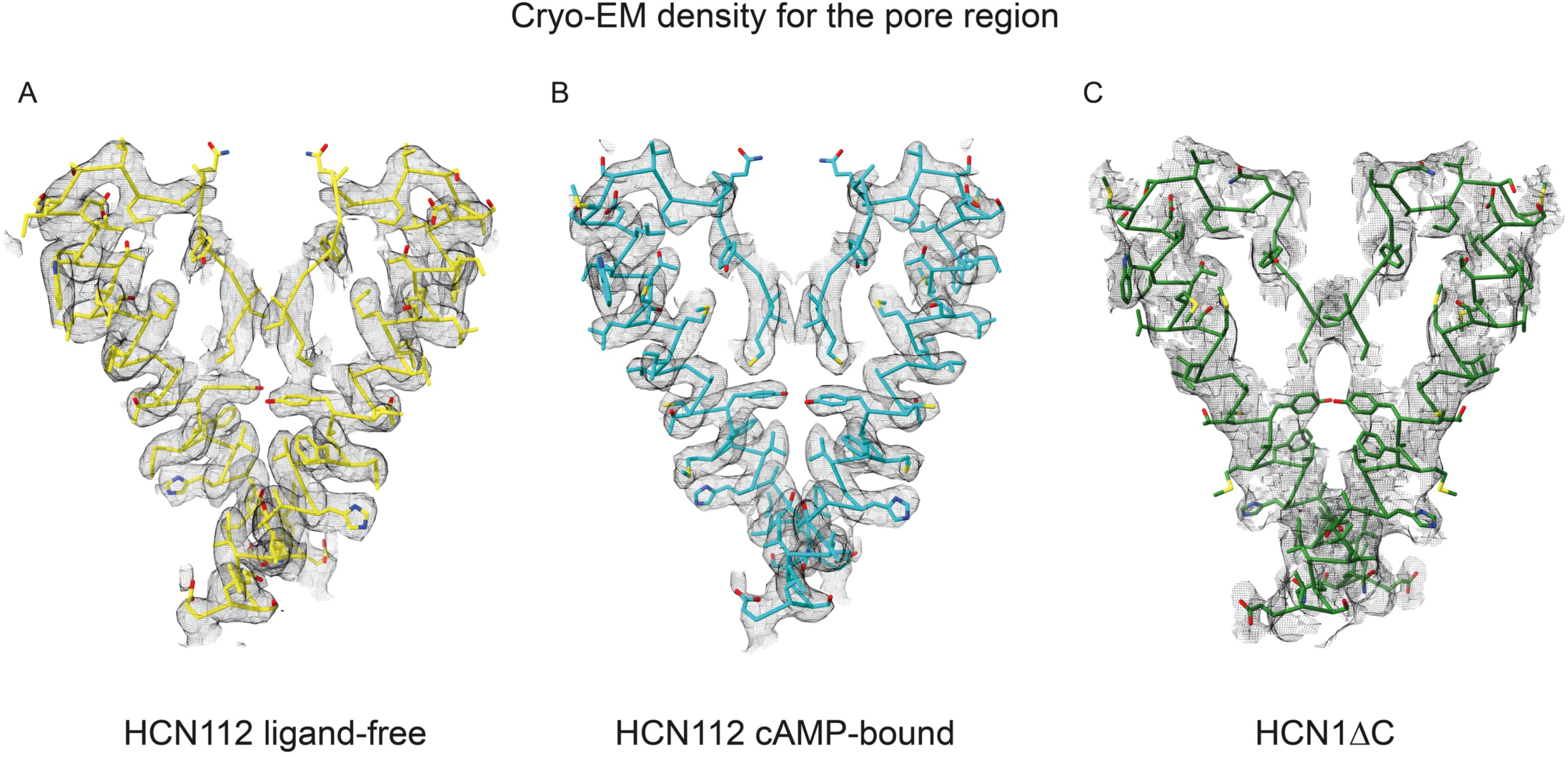
Cryo-EM density for the pore region of HCN112 ligand-free, HCN112 cAMP-bound, and HCN1ΔC. Pore helices (358 to 401 residues) from subunits A and C are shown with their backbones and side chains (in cyan), together with their density (in grey mesh).

**Supplementary Table 1.** Cryo-EM data collection and processing, refinement and validation.

|  | #1 | #2 | #3 |
| --- | --- | --- | --- |
|  | HCN112 ligand-free<br>(EMDB-53516)<br>(PDB 9R1V) | HCN112 cAMP<br>(EMDB-53514)<br>(PDB 9R1T) | HCN1ΔC<br>(EMDB-53515)<br>(PDB 9R1U) |
| **Data collection and processing** |  |  |  |
| Magnification | 96k | 130k | 130k |
| Voltage (kV) | 300 | 300 | 300 |
| Electron exposure (e-/Å<sup>2</sup>) | 42.84 (merge 41.42) | 32.44 | 39.81 |
| Defocus range (μm) | -3.2 to -1.6<br>(0.2 steps) | -3.0 to -1.2<br>(0.2 steps) | -3.4 to -1.6<br>(0.2 steps) |
| Pixel size (Å) | 0.81 | 0.91 | 0.91 |
| Symmetry imposed | C4 | C4 | C4 |
| Initial particle images (no.) | 370859 | 342588 | 1478153 |
| Final particle images (no.) | 92412 | 97137 | 20854 |
| Map resolution (Å) | 3.53 | 3.26 | 3.92 |
| FSC threshold | 0.143 | 0.143 | 0.143 |
| Map resolution range (Å) | 3.53-2.9 | 3.26-2.7 | 3.92-3.45 |
| **Refinement** |  |  |  |
| Initial model used (PDB code) | 5U6O | HCN112 | HCN112 cAMP |
| Model resolution (Å) | 3.5 | 3.3 | 3.1 |
| FSC threshold | 0.143 | 0.143 | 0.143 |
| Model resolution range (Å) | 3.3 | 3.1 | 4.0 |
| Map sharpening *B* factor (Å<sup>2</sup>) | 0.08 | 0.07 | 0.12 |
| Model composition | 4 chains | 4 chains | 4 chains |
| Non-hydrogen atoms | 14668 | 15784 | 8794 |
| Protein residues | 1888 | 2016 | 1148 |
| Ligands | - | 4CMP | - |
| *B* factors (Å<sup>2</sup>) |  |  |  |
| Protein | 61.58 | 75.72 | 187.35 |
| Ligand | - | 93.96 | - |
| R.m.s. deviations |  |  |  |
| Bond lengths (Å) | 0.003 | 0.003 | 0.003 |
| Bond angles (°) | 0.630 | 0.562 | 0.787 |
| **Validation** |  |  |  |
| MolProbity score | 2.06 | 2.08 | 2.74 |
| Clash score | 16.56 | 14.89 | 17.73 |
| Poor rotamers (%) | 0.00 | 1.34 | 4.73 |
| Ramachandran plot |  |  |  |
| Favored (%) | 95.14 | 95.53 | 92.31 |
| Allowed (%) | 4.86 | 4.47 | 7.69 |
| Disallowed (%) | 0.00 | 0.00 | 0.00 |

**Supplementary Movie S1 –** Cartoon representation of 2 opposing HCN112 subunits. The morph illustrates the compression movement of the transmembrane domain versus the cytoplasmic domain. The compression is quantified by measuring the distance between the Calpha atom of residue Q323 at the top of the S5 helix and residue G547 at the distal end of the CNBD (indicated as yellow spheres). The compression amounts to ∼3.5 Å.

**Supplementary Movie S2** – Conformational change of the CNBD upon binding of cAMP in the HCN112 channel. 2 opposing subunits are shown in cartoon representation.

**Supplementary Movie S3** – Conformational change of the transmembrane domain (especially S5 and S6) relative to the HCND domain. 2 opposing subunits of HCN1ΔC are shown in cartoon representation. mental Videos and Spreadsheets

## CELL PRESS DECLARATION OF INTERESTS FORM

If submitting materials via Editorial Manager, please complete this form and upload with your initial submission. Otherwise, please email as an attachment to the editor handling your manuscript.

*Please complete each section of the form and insert any necessary “declaration of interests” statement in the text box at the end of the form. A matching statement should be included in a “declaration of interests” section in the manuscript*.

### Institutional affiliations

We require that you list the current institutional affiliations of all authors, including academic, corporate, and industrial, on the title page of the manuscript. ***Please select one of the following:***

☐ All affiliations are listed on the title page of the manuscript.

☐ I or other authors have additional affiliations that we have noted in the “declaration of interests” section of the manuscript and on this form below.

### Competing financial interests

We require that authors disclose any financial interests and any such interests of immediate family members, including financial holdings, professional affiliations, advisory positions, board memberships, receipt of consulting fees, etc., that:

1. could affect or have the perception of affecting the author’s objectivity, *or*
2. could influence or have the perception of influencing the content of the article.

***Please select one of the following:***

☐ We, the authors and our immediate family members, have no financial interests to declare.

☐ We, the authors, have noted any financial interests in the “declaration of interests” section of the manuscript and on this form below, and we have noted interests of our immediate family members.

### Advisory/management and consulting positions

We require that authors disclose any position, be it a member of a board or advisory committee or a paid consultant, that they have been involved with that is related to this study. We also require that members of our journal advisory boards disclose their position when publishing in that journal. ***Please select one of the following:***

☐ We, the authors and our immediate family members, have no positions to declare and are not members of the journal’s advisory board.

☐ The authors and/or their immediate family members have management/advisory or consulting relationships noted in the “declaration of interests” section of the manuscript and on this form below.

### Patents

We require that you disclose any patent applications and/or registrations related to this work by any of the authors or their institutions*. **Please select one of the following:***

☐ We, the authors and our immediate family members, have no related patent applications or registrations to declare.

☐ We, the authors, have a patent application and/or registration related to this work, which is noted in the “declaration of interests” section of the manuscript and on this form below, and we have noted the patents of immediate family members.

***Please insert any “declaration of interests” statements in this space.*** This exact text should also be included in the “declaration of interests” section of the manuscript. If no authors have a competing interest, please insert the text, “The authors declare no competing interests.”

- ☐ **On behalf of all authors, I declare that I have disclosed all competing interests related to this work. If any exist, they have been included in the “declaration of interests” section of the manuscript.**

